# Toward a Minimal Amino Acid Alphabet for Protein Design

**DOI:** 10.64898/2026.03.06.710107

**Authors:** Karel Půbal, Kseniia Kushnir, Vojtěch Spiwok, Karolína Loužecká, Vladimír Setnička, Petra Lipovová

## Abstract

Proteins are built from 20 canonical amino acids. It is interesting to explore whether proteins can be formed from significantly reduced amino acid alphabets. Our bioinformatics survey of UniProt (more than 250 M sequences) revealed that proteins composed of reduced amino acid alphabets (< 10) are extremely rare among existing proteins. Next, we used computational protein design to design proteins composed of all 1,013 possible alphabets of 2-10 early amino acids (Ala, Asp, Glu, Gly, Ile, Leu, Pro, Ser, Thr, and Val). The length of all proteins was 100 amino acid residues. Small amino acid alphabets preferred simple helices or helix bundles. Larger amino acid alphabets allowed for the design of more complex structures. A protein composed of 8 amino acid types (Ala, Asp, Gly, Leu, Val, Ser, Thr, and Pro) was successfully experimentally verified. It adopts the β-sheet-rich fibronectin type III domain architecture. Attempts to experimentally verify designs composed of 6 and 4 amino acid types were unsuccessful. We show by a computational experiment with an experimental validation that inverse folding models, namely ProteinMPNN_sol_, can stabilize a designed protein within the same eight-amino-acid alphabet. Our results show that globular proteins may have formed early in evolution. Furthermore, we show that it is possible to design proteins with interesting properties for biotechnology and synthetic biology.

## 1 Introduction

Proteins can perform different functions including signaling, building, catalytic, and many others. The ability to form well-defined 3D structures is essential for many of these functions. The question arises whether structured proteins can be formed from the number of amino acids significantly lower than in the modern 20-amino-acid alphabet.

This question is interesting from an evolutionary point of view (Longo and Blaber (2012); Higgs and Pudritz (2009)). It is believed that 20 amino acids can be divided into early and late types. The early amino acids (Ala, Asp, Glu, Gly, Ile, Leu, Pro, Ser, Thr, and Val) are supposed to be present on pre-biotic Earth (approximately 5 GA). There is a great deal of evidence, for example, the Miller-Urey experiment (Miller (1953)), experiments simulating hydrothermal vents (Huber and Wächtershäuser (2006)) and the analysis of carbonaceous meteorites (Cronin and Moore (1971); Shimoyama et al. (1979); Engel and Nagy (1982)), that early amino acids can be formed abiotically. In contrast, late amino acids (Cys, Phe, His, Lys, Met, Asn, Gln, Arg, Trp, and Tyr) are supposed to emerge from the metabolic pathways of early organisms. Some studies propose that sulfur amino acids (Van Trump and Miller (1972); Parker et al. (2011)) or histidine (Shen et al. (1990)) could also have been present in the pre-biotic environment; nevertheless, we will use the term “early amino acids” only for the set of 10 amino acids referred to above.

It is evident that the proteins of the early organisms must have been built from a reduced amino acid alphabet, most likely from early amino acids. What did such proteins look like? Were they soluble globular proteins, similar to modern ones? Or some amyloid-like structures? Or quasi-random coacervates? We hypothesize that such proteins included fully folded globular proteins. To address this question, one must ask whether the reduced amino acid alphabet is large enough to form globular proteins.

The question is interesting from the biotechnological point of view as well. Proteins built from reduced amino acid alphabets may be more difficult to recognize for proteolytic enzymes and other degrading mechanisms. This can increase their stability in biological systems. Similarly, reduced-alphabet proteins can be recognized by the immune system differently from full-alphabet proteins. All of these properties would be interesting for therapeutic and other biotechnological applications.

Numerous studies have attempted to develop proteins from minimal amino acid alphabets. Schafmeister and co-workers (Schafmeister et al. (1997)) designed a protein composed of 7 amino acid types using a computer-graphics-guided protein design. This protein folds into a four-helix bundle and is one of the first *de novo* designed proteins. Another computationally designed protein is one of two-state switchable hinge proteins by Praetorius and co-workers (PDB ID 8FVT) (Praetorius et al. (2023)) from ten amino acid types (namely AEGHKLRSTV). To the best of our knowledge, this study did not focus on amino acid alphabet size.

Several low-alphabet proteins have been developed by a reduction of the alphabets of selected existing (not designed) proteins. Riddle and co-workers (Riddle et al. (1997)) selected the SH3 domain (108 residues) as the model system. They retained amino acid residues essential for its biological function and replaced those responsible for the formation of the 3D structure. The resulting proteins composed of as few as 5 amino acid types (plus those essential for function) retained their activity, which is the molecular recognition of partner proteins. Longo, Lee, and Blaber (Longo et al. (2013)) developed variants of β-trefoil proteins composed of 12 or 13 residues, stable at high salt concentrations. Walter, Vamvaca, and Hilvert (Walter et al. (2005)) developed variants of chorismate mutase composed of 9 amino acid types that retain their enzymatic activity. Shibue and co-workers (Shibue et al. (2018)) systematically reduced the amino acid alphabet of nucleoside kinase fold down to 13 amino acids, yielding soluble, stable, and catalytically active proteins.

Ancestral sequence reconstruction, Rosetta scoring, molecular simulations, and other approaches were used by Yagi and co-workers (Yagi et al. (2021)) to design double-ψ β-barrel proteins from seven amino acid types. The amino acid types they used can be coded by GNN and ARR (N = any nucleotide, R = A or G) codons, which implies that such a protein could be produced by early translation systems and genetic codes. Makarov and co-workers (Makarov et al. (2023)) systematically tested solubility and other properties of peptides composed of pre-biotic types of amino acids (both coded and non-coded). They demonstrated that foldability was a selective pressure that likely led to the formation of an early alphabet of ten amino acids and subsequently into the modern amino acid alphabet.

Another insight comes from Especial and Faísca (Especial and Faísca (2025)) who studied reduced amino acid alphabets in the context of protein knots. They used Gō model and Monte Carlo sequence sampling to redesign three proteins that differ in the degree of knotting. It was shown that the early amino acid alphabet disfavors deep knots in proteins.

Recently, Giacobelli and co-workers (Giacobelli et al. (2025)) used artificial-intelligence-driven protein design, namely RFdiffusion (Watson et al. (2023)) in the fold conditioning mode to redesign selected evolutionarily old proteins (DNA/RNA-binding 3-helical bundle, ferredoxin, and RNase H). Next, they used Protein Message Passing Neural Network (Dauparas et al. (2022)), to provide amino acid sequences for these folds composed exclusively from 10 early amino acids. The selected redesigned variants were prepared and structurally characterized, and it was demonstrated that they are highly thermostable.

Finally, proteins composed of reduced amino acid alphabets have been intensively studied using theoretical methods (Fan and Wang (2003); Nerattini et al. (2020)).

The purpose of this study is to test how many amino acid types are needed to build a globular protein. First, we searched for such proteins among modern proteins in UniProt. Next, we attempted to computationally design and experimentally evaluate such proteins. Finally, we used an inverse folding model to further stabilize one of the designed proteins.

## 2 Results

### 2.1 Bioinformatics survey on modern proteins

As a first step, we investigated how many modern proteins are built from reduced amino acid alphabets (Figure 1A). We started from UniProt (sequences of 253,061,697 proteins) (The UniProt Consortium (2024)) and analyzed them to find proteins built from fewer than 10 amino acid types and having at least 100 amino acid residues. This resulted in 29,457 proteins, which is 0.012 % of all proteins. We also removed proteins larger than 1,000 amino acid residues and those with unknown amino acids to enable structure prediction using ESMfold (Lin et al. (2023)). This resulted in 28,843 proteins (0.011 %). Next, we retrieved those with ESMfold pLDDT greater than 80. This led to 2,794 proteins.

**Figure 1.**
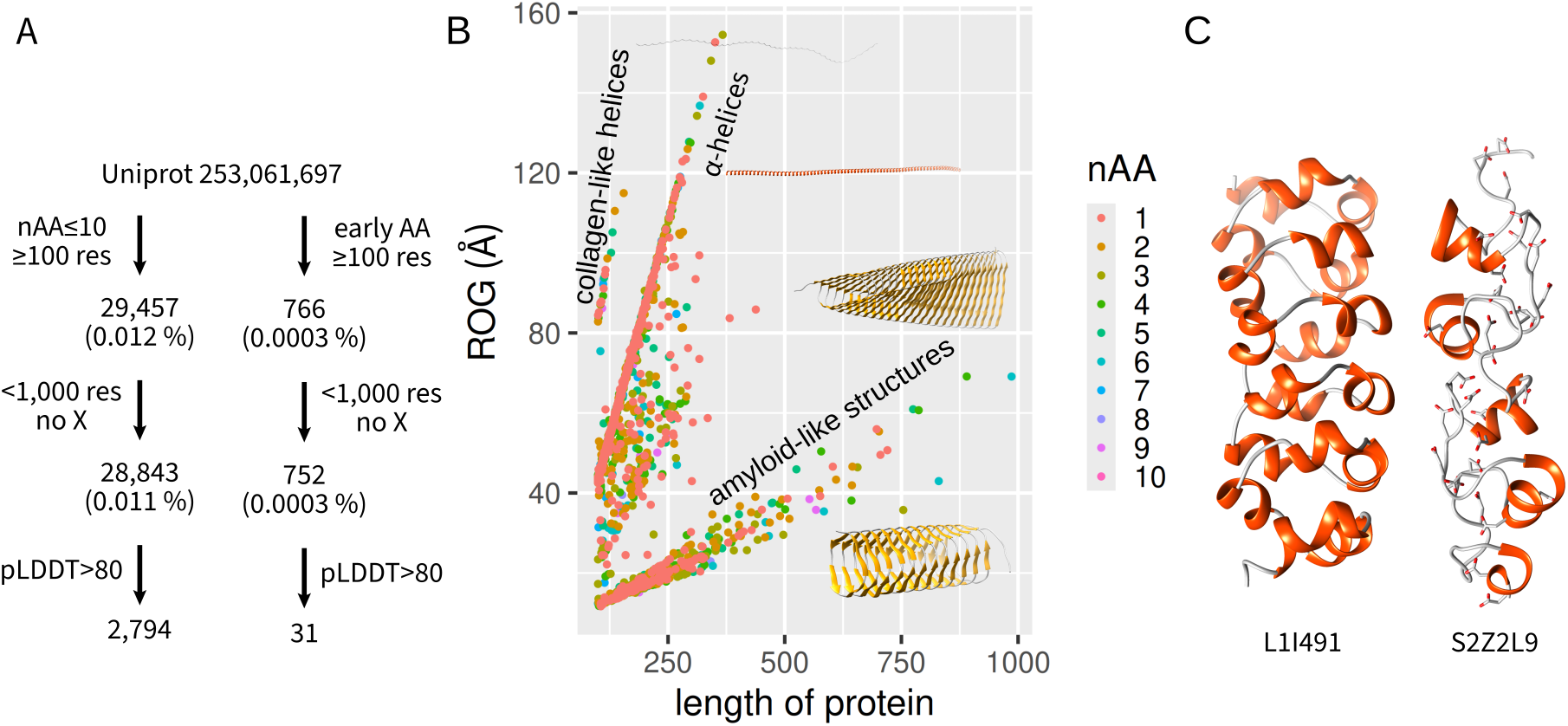
Utilization of reduced amino acid alphabets by modern proteins. (A) Workflow for finding modern proteins composed of any reduced amino acid alphabet (left) or a reduced alphabet from early amino acids (right). (B) Analysis of the resulting 2,794 proteins. Each protein is represented as a dot colored by the number of amino acid types from which it is formed. Lines pointing from the origin correspond to repetitive structures. (C) Selected interesting proteins L1I491 (uncharacterized protein from *Guillardia theta*) and S2Z2L9 thrombospondin type 3 repeat-containing protein of *Corynebacterium pyruviciproducens*.

They are analyzed in Figure 1B. Each protein is a dot on a scatter plot with the number of amino acid residues on the horizontal axis and the radius of gyration on the vertical axis. Proteins with highly repetitive structures form lines that point to the origin. This is the case for collagen-like helices, long α-helices, or amyloid-like structures.

Further analysis confirmed that there were no α/β-proteins (except for some amyloid-like structures with a short terminal helix). We also focused on proteins with multiple α-helices and found very few structurally interesting examples, such as those depicted in Figure 1C.

We highlight proteins with the UniProt IDs L1I491 (uncharacterized protein from *Guillardia theta*, Curtis et al. (2012)) and S2Z2L9 (thrombospondin type 3 repeat-containing protein of *Corynebacterium pyruviciproducens*, unpublished). Both proteins contain repetitive structural motifs. The latter is probably a metal-binding protein, as indicated by the number of aspartic acid residues that point to the center (depicted in Figure 1C).

A similar analysis was performed for early amino acids (instead of all alphabets of size 1-10 formed by any amino acid types). There were 766 proteins (≥100 residues) composed of early amino acids. Thirty-one of them had an ESMfold pLDDT greater than 80 and none of them were identified as globular proteins.

In the Protein Databank (PDB), we found two previously mentioned designed proteins (PDB IDs 4HB1 and 8FVT) (Schafmeister et al. (1997); Praetorius et al. (2023)), amyloid-like fibrils formed by a low-complexity region from the FUS protein (PDB ID 6XFM) (Lee et al. (2020)) and collagen structures (PDB IDs 8ZXL and 9IUS, unpublished) as only representative of proteins composed of ten or fewer amino acid types.

In conclusion, we can say that reduced amino acid alphabets are not used in modern proteins and that modern proteins composed of reduced amino acid alphabets are extremely rare and mostly non-globular. In general, nature now uses the full amino acid alphabet. We admit that there could be some limitations in this survey, for example, in structure prediction by ESMfold or due to the fact that we omitted proteins longer than 1,000 amino acid residues. However, we believe that this does not change the overall conclusion.

#### 2.1.1. Design of proteins with reduced amino acid alphabets

As the next step, to test whether proteins composed of reduced amino acid alphabets can fold into compact structures, we attempted to design them using computational protein design. We used a modified version of the ESMfold-based module known as language model design (`lm-design`, Verkuil et al. (2022)). This program uses a joint sequence-structure probability to score an amino acid sequence for its propensity to form a compact 3D structure. The design starts from a random amino acid sequence, which is optimized by Monte Carlo simulated annealing. We modified this program in order to enable work with reduced amino acid alphabets and to add additional output.

Initially, we wanted to analyze all possible amino acid alphabets. It turned out that the number of combinations for 2-20 amino acid types is 1,048,584, which exceeds our computational resources. Restriction to all amino acid alphabets of size 2-10 reduces the number of combinations to 616,646, which is still not feasible. Therefore, we decided to focus on 10 early amino acids. The total number of possible amino acid alphabets of size 2-10 formed only by early amino acids is 1,013. This corresponds to one alphabet of all early amino acids, 10 alphabets of 9 amino acid types, 45 alphabets of 8 amino acid types, etc. (Figure 2A).

**Figure 2.**
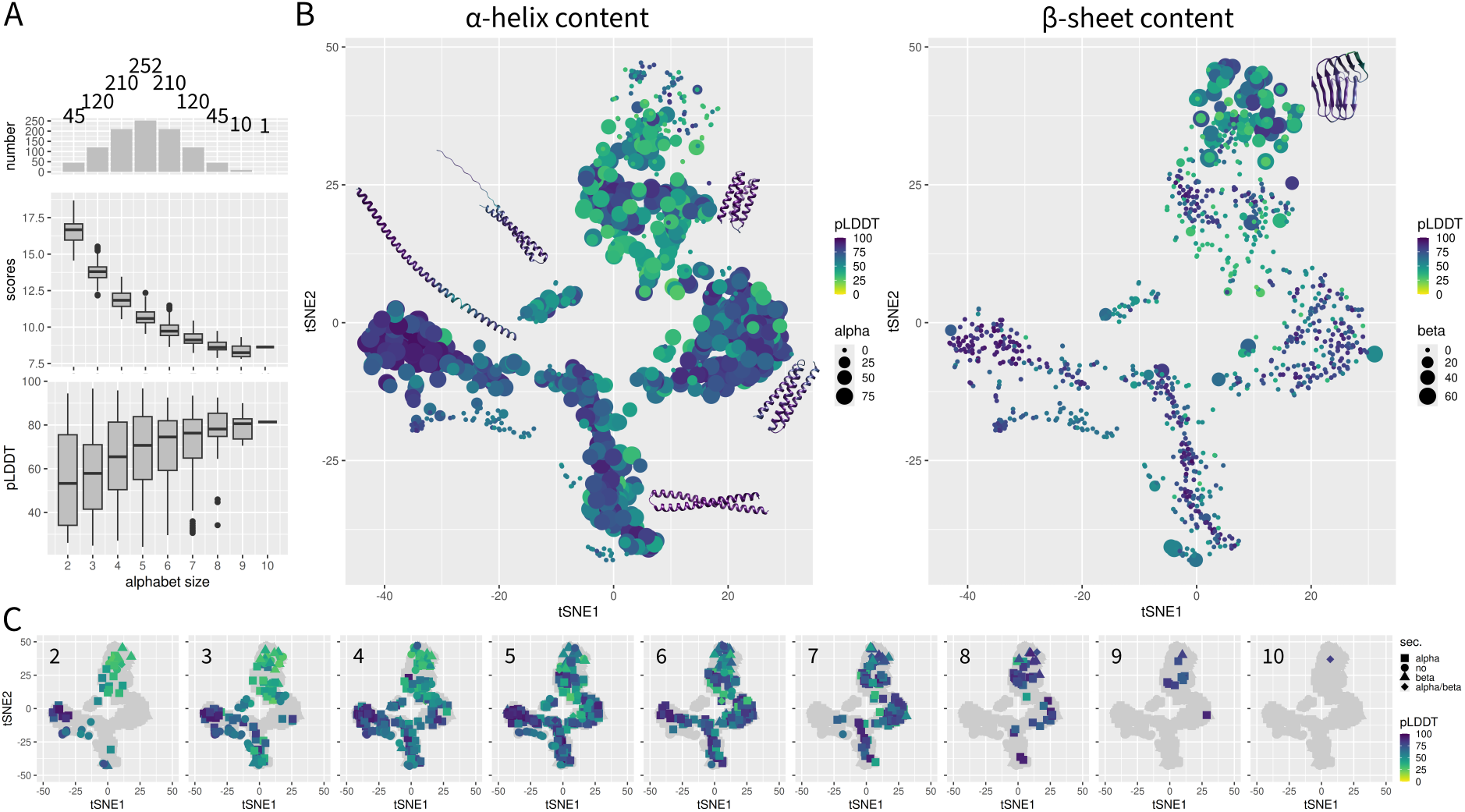
Design of proteins from reduced amino acid alphabets. (A) Numbers of combinations (top), final scores (middle), and final ESMfold pLDDT scores for alphabets of different sizes (2-10). (B) tSNE analysis of structures of designed proteins. Each designed protein is depicted as a point. Its color indicates pLDDT and the size indicates α-helix (left) or β-sheet (right) content. Representative structures are also depicted (colored by pLDDT with the same scale). (C) tSNE plots for individual alphabet sizes. Colors and shapes of points indicate pLDDT and protein type, respectively (models from other alphabet sizes are in gray).

We computationally designed one protein for each of 1,013 amino acid alphabets. Each designed protein had exactly 100 amino acid residues. The 3D structure of each designed protein was predicted using ESMfold. The secondary structure was identified by DSSP (Hekkelman et al. (2025)). Proteins were classified as α, β or α/β if they contained any content of the respective secondary structure type.

Figure 2A shows the number of combinations, scores, and pLDDT values of the final models. The score decreases with the size of the alphabet. This can be explained by its probabilistic nature rather than by the quality of the designed sequences, since the probability of a certain sequence depends on the size of the alphabet. We designed high-pLDDT models for all alphabet sizes. High-pLDDT models were prevalent for alphabets of sizes 8, 9, and 10.

To compare 3D structures, each structure was aligned with the protein design of all 10 early amino acids (which was used as a reference) and the backbone coordinates were analyzed using tSNE (van der Maaten and Hinton (2008)). The alignment step and the tSNE analysis did not take into account the amino acid sequence. The results are depicted in Figure 2B. The tSNE plot shows multiple clusters corresponding to different types of structure. They include fully unstructured proteins, partially or fully structured long α-helices, two-helix and three-helix bundles, and more complex structures. More complex structures included α/β-proteins, β-rich proteins, including amyloid-like structures, and helix bundles. There was a clear trend for very small alphabets to form unstructured proteins or proteins with non-complex structures.

Figure 2C further separates amino acid alphabets of different sizes. Small amino acid alphabets (2-5) predominantly form low-complexity structures such as long helices or two-helix bundles. Larger alphabets provide more complex designs. A large set of designed proteins allowed us to elucidate the effect of the presence of amino acid types on protein designability. The score optimized in `lm-design` or the pLDDT of the ESMfold-predicted model were used as estimators of the designability. The stabilizing (pro-design) amino acid types were those whose presence in the alphabet tends to decrease the score or increase pLDDT. We constructed a regression model of the score or pLDDT as a function of the size of the alphabet and the presence or absence of individual amino acid types.

Figure 2A clearly shows that the scores and pLDDT values strongly depend on the size of the alphabet. Therefore, we used a simple linear model *score* = *an* + *b* and *pLDDT* = *a*^*′*^*n* + *b*^*′*^, where *n* is the size of the alphabet and *a, a*^*′*^, *b* and *b*^*′*^ are parameters of models. The residuals of these models were then fitted by models *residuals* = *α*_*A*_*A* + *α*_*D*_*D* + …, where *A* is 1 if Alanine is present, 0 if absent, etc. for all early amino acids. This allowed us to remove the effect of alphabet size and focus on the effects of presence of individual amino acids. These effects were expressed as the shift of the score (positive shift means a destabilizing effect and *vice versa*) and the shift of the pLDDT (positive shift means a stabilizing effect and *vice versa*). Figure 3 shows the effects of individual amino acids on designability. Serine was identified as a neutral amino acid with an insignificant effect (*P >* 0.05). A significant destabilizing effect (*P <* 0.001) was observed for Glycine and Proline, both for score and pLDDT. A significant stabilizing effect was observed for Isoleucine, Alanine, and Glutamic acid.

**Figure 3.**
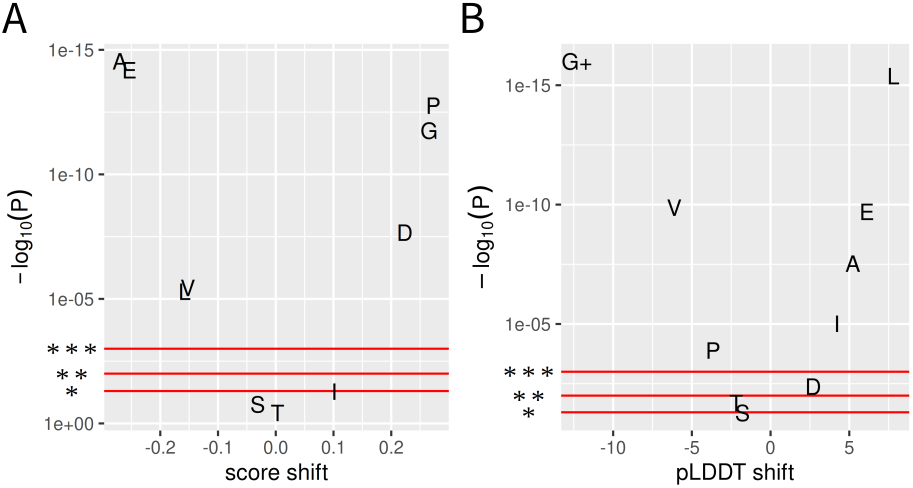
Effect of individual amino acid types on the designability. The effect of the presence of individual amino acid was evaluated as the trend of decreasing the lm-design score or increasing of pLDDT by the presence of the amino acid in the alphabet. (A) The effect on score (positive means destabilizing). (B) The effect on pLDDT (positive means stabilizing, + *P*-value of G was *<* 2.2 · 10^−16^); * *P* = 0.05; ** *P* = 0.01; *** *P* = 0.001

Since knots in protein structures attracted attention from an evolutionary point of view (Especial and Faísca (2025)), we analyzed all 1,013 proteins to determine the presence of knotted structures. We observed six proteins with a probability of a 3_1_ knot greater than 0.5. However, visual inspection revealed that all these knots were present in the low-pLDDT parts, so they are likely artifacts of 3D structure prediction.

### 2.2 Experimental validation

Three designs composed of 8, 6, and 4 amino acid types were selected for experimental validation. This selection was driven by the need to cover different sizes of amino acid alphabets. We selected proteins with low lm-design scores (relative to other designs of the same alphabet size) and high pLDDT values. Visual inspection of predicted 3D structures was also used in the selection process.

The selected designs were recombinantly expressed in *E. coli* in a fusion with the His-tag and thrombin cleavage site (Table 1). The designs were then isolated by immobilized metal affinity chromatography followed by gel permeation chromatography (GPC). The purpose of the second chromatography was not only to purify the protein but also to assess its shape from its retention time. The successfully purified proteins were analyzed by electronic circular dichroism (ECD). The His-tag was not removed for simplicity.

**Table 1.**
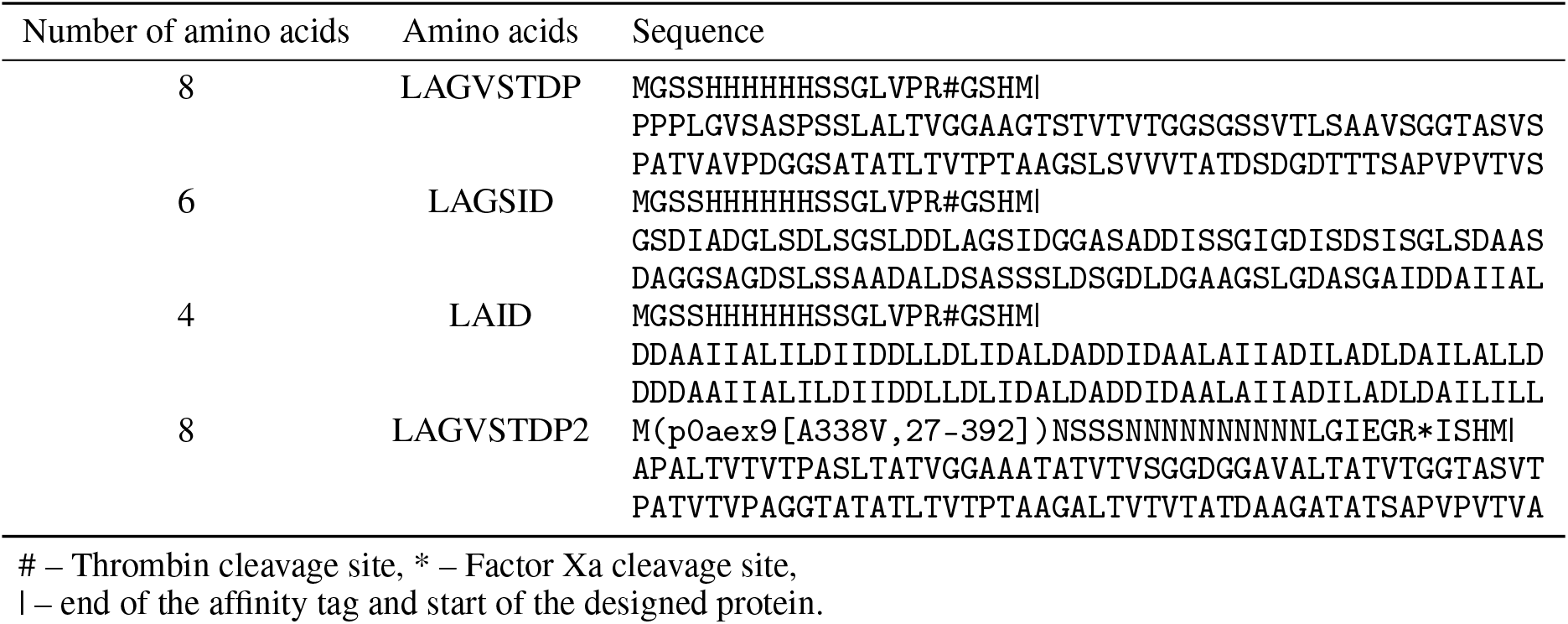
Experimentally validated protein constructs (maltose binding protein sequence is replaced by the corresponding UniProt ID, mutation and sequence range).

The first protein tested experimentally was LAGVSTDP (Figure 4) formed of 8 amino acid types (Leu, Ala, Gly, Val, Ser, Thr, Asp, and Pro). The resulting score was 9.74 and the pLDDT was 91.6. It was predicted to fold into the fibronectin type III domain, as identified by FoldSeek (van Kempen et al. (2024)) (Figure 4A). The model shows strong 3D similarity to a domain of the tomato subtilisin-like protease SBT3 (PDB ID 3I6S, Ottmann et al. (2009)). In terms of sequence, it shows 47 % identity with metallophosphoesterase from *Stackebrandtia nassauensis* (UniProt ID D3Q2J9, Aravind and Koonin (1998)). Note that both proteins use almost complete amino acid alphabets (the former without Cys and Met and the latter without Cys, Met, and Trp).

**Figure 4.**
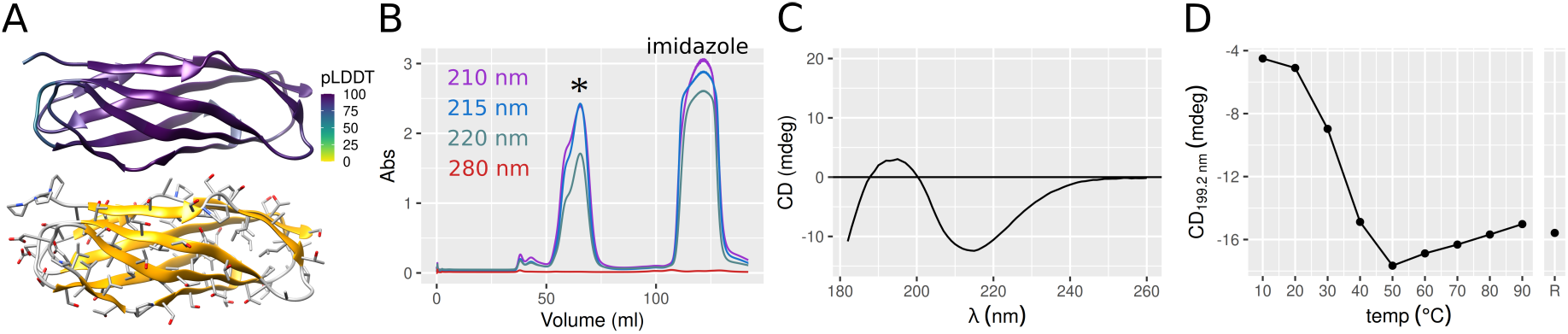
Experimental testing of LAGVSTDP. (A) Predicted 3D structure colored by pLDDT (top) or secondary structure/atom type (bottom). (B) Gel permeation chromatography elution profile (* indicates LAGVSTDP). (C) Electronic circular dichroism spectrum of LAGVSTDP. (D) Temperature denaturation profile of LAGVSTDP. R refers to a refolding attempt (cooling back to 20 °C after 90 °C).

LAGVSTDP was successfully expressed in *E. coli* and separated by metal affinity chromatography and GPC. It was eluted from GPC as a second of two partially overlapping peaks (Figure 4B). It was demonstrated that LAGVSTDP, once separated from the protein that forms the first peak, remains stable and does not convert into the protein that forms the first peak (see the Zenodo dataset). The elution volume of LAGVSTDP supports its proper folding.

This was further confirmed by the ECD spectrum (Figure 4C). The shape of the ECD spectrum corresponds to a β-sheet-rich protein. A thermal unfolding experiment (Figure 4D) showed that the protein is stable only at low temperatures with melting temperature between 30 and 40 °C. The unfolding was irreversible.

We attempted to recombinantly produce proteins composed of 6 (LAGSID from Leu, Ala, Gly, Ser, Ile, and Asp) and 4 (LAID, from Leu, Ala, Ile, and Asp) amino acid types. Unfortunately, we were unable to obtain enough protein to characterize it by ECD. Although a low level of expression in *E. coli* was detectable by Western blot, the amount of protein produced was insufficient for a successful characterization.

### 2.3 Redesign with experimental validation

The designed protein LAGVSTDP was poorly thermostable (Figure 4D). We attempted to redesign this protein using Protein Message Passing Neural Network trained on soluble proteins (ProteinMPNN_sol_, Goverde et al. (2024)) using the same amino acid types (Figure 5A). The ESMfold model of LAGVSTDP was used as input for ProteinMPNN_sol_ and 50 sequences were generated. The redesigned sequences (one with the highest confidence), further referred to as LAGVSTDP2, formed the same 3D structure according to the ESMfold. Its confidence was higher than for the original LAGVSTDP (0.56 for LAGVSTDP2 vs. 0.39 for LAGVSTDP). This indicates that the low stability of LAGVSTDP can be explained by the limitations of `lm-design`. Similar results (i.e. improvement of confidence and the retention of the 3D structure according to ESMfold) were observed for LAGSID and LAID, however, their confidence was much lower.

**Figure 5.**
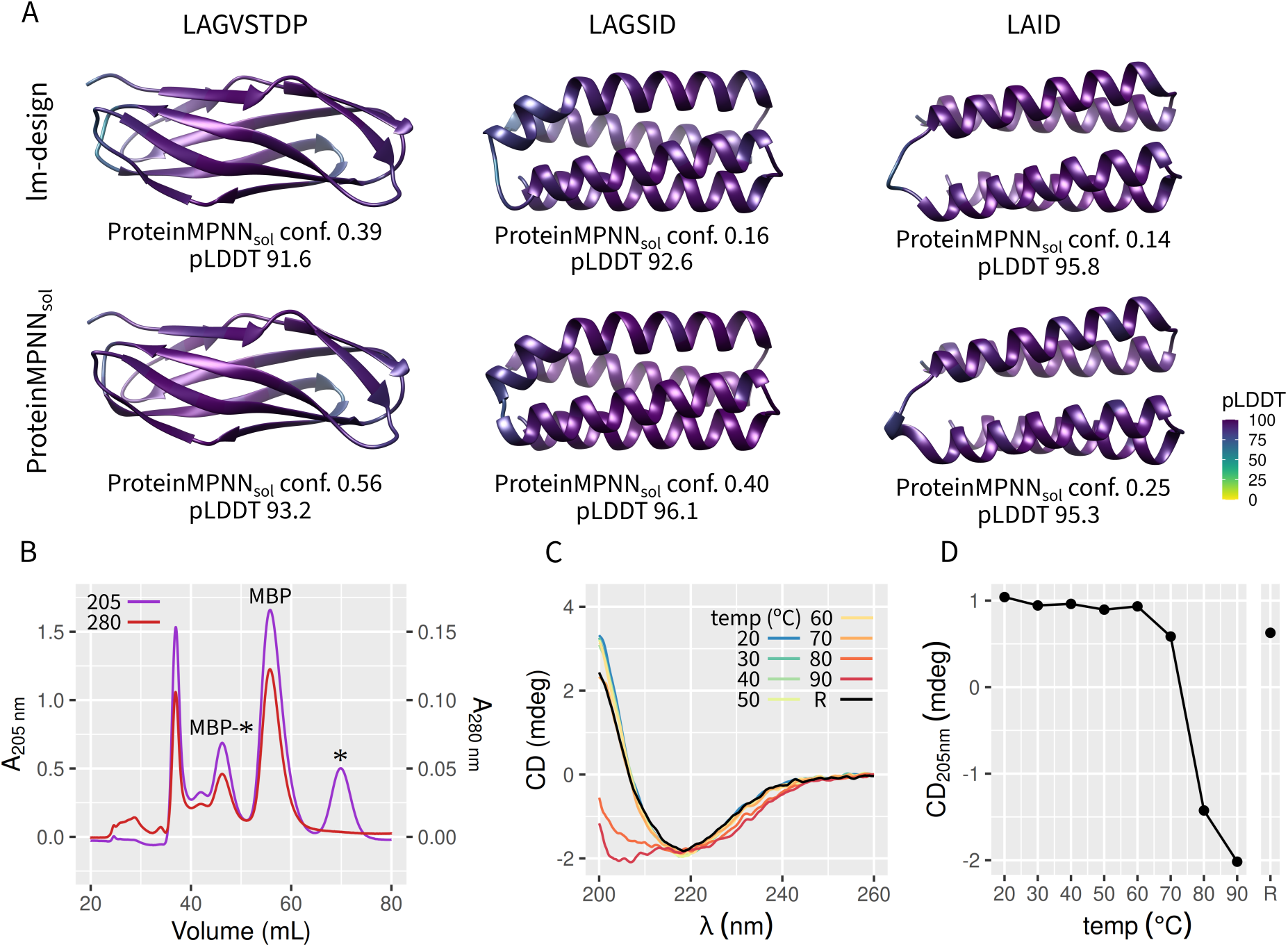
Redesign using ProteinMPNN_sol_. (A) The proteins LAGVSTDP, LAGSID and LAID were redesigned by ProteinMPNN_sol_. The figures on the top are ESMfold models designed by lm-design, the figures on the bottom were redesigned by ProteinMPNN_sol_. The figures are colored by pLDDT. The values of ProteinMPNN_sol_ confidences and average pLDDT are also indicated. (B) Gel permeation chromatography elution profile (* indicates LAGVSTDP2, MBP indicates maltose-binding protein). (C) Electronic circular dichroism spectrum of LAGVSTDP2 at different temperatures. (D) The temperature denaturation profile of LAGVSTDP2. R refers to a refolding attempt (cooling back to 20 °C after 90 °C).

The sequence identity of LAGVSTDP2 with LAGVSTDP was 69.7 %. LAGVSTDP2 was similar, in terms of sequence, to the calcineurin-like phosphoesterase domain-containing protein of *Allocatelliglobosispora scoriae*. The ESMfold model of LAGVSTDP2 was most similar (33 % sequence identity) to the *Marinomonas primoryensis* peptide-binding domain (PDB ID: 6X6M, Guo et al. (2021)). Unmodeled parts of other proteins in PDB were not considered.

The redesigned variant of LAGVSTDP (LAGVSTDP2, Table 1) was experimentally verified. In order to improve the process of recombinant protein expression, we replaced a His-tag by maltose binding protein (MBP). MBP was cleaved by factor Xa prior to the ECD measurement (Figure 5B). ECD revealed that the LAGVSTDP2 was thermostable (Figure 5C,D). It was stable up to 70 °C with melting temperature between 70 and 80 °C. More importantly, the unfolding was reversible, as demonstrated by the recovery of the ECD spectrum back to the original profile after cooling to 20 °C.

## 3 Discussion

We have not found any interesting natural (not designed) proteins composed of small amino acid alphabets in our bioinformatics survey of UniProt. We must admit that there could be some limitations in structure prediction using ESMfold, namely that some proteins predicted as unstructured are, in fact, structured. Low values of pLDDT could mean that a protein or its region is not structured or that ESMfold or AlphaFold does not properly model it. Nevertheless, since some amyloid-like, helical, or collagen-like proteins were modeled correctly, we believe that the overall conclusion that modern proteins use the full amino acid alphabet and small-alphabet proteins are extremely rare is correct.

An interesting fact is that many natural proteins composed of small amino acid alphabets are amyloid-like, i.e., highly periodic structures with a high content of *β*-sheets. In addition, numerous amyloid-like structures were devised in our design campaign. This indicates that amyloid-like structures might have played an important role in early evolution. Today, amyloids are perceived as a large group of proteins with many functions, not only in the context of misfolding pathologies (Golan et al. (2022)). From a protein design perspective, the presence of amyloid-like structures among designed proteins may be explained by the fact that with a reduced amino acid alphabet, the program seeks inspiration in existing low-amino-acid-alphabet proteins.

In the *de novo* designed proteins, we observed a clear relationship between the structure and alphabet size. Figure 2C shows that alphabets of sizes 2-5 lead to long α-helices (left part of the tSNE plot) with high pLDDT, helical bundles, and compact but poorly structured designs at the top of the tSNE plot. Moderately sized alphabets form predominantly helical bundles. Complex protein structures were obtained for alphabets of size 8-10. We did not observe any signs of preferential presence of knots in designed structures, at least in the globular parts.

Helical structures dominated the designed proteins, especially those from small alphabets. In general, helical proteins are recognized as easier to design than β-sheet-rich proteins (Koh et al. (2025)). Therefore, one possible explanation is that the design method fails to produce β-sheet-rich proteins in difficult design tasks. However, we can speculate that this is due to the propensity of small amino acid alphabet proteins, rather than a limitation of the protein design method.

Figure 2A shows that the final scores of `lm-design` regularly decrease with increasing alphabet size. The intervals for individual alphabet sizes are very narrow. In contrast, pLDDT forms wide intervals. The ranges of these intervals decrease with the size of the alphabet. We can explain this difference by the fact that the score is defined by the probabilities of the presence of different residues in different positions. The probability and thus the score are biased by the alphabet size.

We observed some trends in which some amino acid types supported and others hindered protein design (Figure 3). These trends are in agreement with the current knowledge of the stabilizing and destabilizing effects of amino acids on designed proteins (Rocklin et al. (2017)). This agrees with the fact that ProteinMPNN penalizes glycine residues in secondary structure elements (Dauparas et al. (2022)).

Reducing the amino acid alphabet poses a challenge for protein language models. A smaller number of building blocks simplifies tasks related to structural analysis and prediction, but at the cost of limiting the information available for training. This was studied by Ieremie and co-workers (Ieremie et al. (2024)). They trained protein language models by sequences in which some groups of amino acid types were represented by one letter. These models performed well for small or moderate reduction and their performance dropped sharply for sequences represented by less than 10 symbols. Interestingly, this is a trend similar to our results. In our work, we reduced the amino acid alphabet by the withdrawal of amino acid types, Ieremie and co-workers reduced alphabets by grouping amino acid types.

The study used the `lm-design` ESMfold module for protein design. The structures of the designed sequences were predicted by ESMfold. The fact that the same approach was used to design proteins and predict their structures may be a limiting factor. Therefore, we predicted the structures of all proteins in Table 1 using AlphaFold 2 (Jumper et al. (2021)). The predicted structures were nearly identical to those predicted by ESMfold (see the Zenodo dataset).

Recently, Giacobelli and co-workers (Giacobelli et al. (2025)) used a computational protein design to demonstrate that globular proteins can be formed of early amino acids. They used a targeted design instead of a free design used in this work. They selected nine folds that have been proven as ancient (present in early organisms) in the literature. Then, they used RFDiffusion (Watson et al. (2023)) and Protein Message Passing Neural Network (ProteinMPNN, Dauparas et al. (2022)) to redesign sequences so that they contain only early amino acids. The results were successfully experimentally validated for three folds using recombinant expression, ECD, and NMR.

The proteins designed by Giacobelli and co-workers (Giacobelli et al. (2025)) were highly thermostable, in contrast to the LAGVSTDP designed in this study. This can be explained by the software used. ProteinMPNN is known to produce highly thermostable proteins and can be used to redesign proteins to increase thermostability. To the best of our knowledge, the thermostability of proteins designed using `lm-design` has not been systematically experimentally tested.

It is appealing but wrong to explain the fact that proteins from early amino acids designed by ProteinMPNN are thermostable by high temperatures on Earth during early evolution. We agree with the explanation of Giacobelli and co-workers that the thermostability can be explained by the method used.

Our study used a free design of proteins (3D structures are being designed together with sequences), whereas Giacobelli and co-workers chose targeted design (3D structures are used as input and only sequences are designed). The advantage of free design in the search for a minimal amino acid alphabet lies in the fact that the 3D structure can, at least in principle, adapt to the alphabet. We can speculate that only some folds can be formed by small amino acid alphabets and free design can search for them. Alternatively, it is possible to collect a large library of folds, either natural or *de novo* designed, and to test each alphabet on the whole library by targeted design, e.g. by ProteinMPNN.

We admit that the designability of proteins from reduced amino acid alphabets may be influenced by many factors, such as the approach (free vs. targeted design), the program, size of proteins, and other details. A detailed analysis would be required to determine whether `lm-design` reuses motifs from existing proteins with low amino acid complexity, or “hallucinates” new motifs suitable for reduced amino acid alphabets.

Our design campaign used a fixed protein length of 100 amino acids. This can be a limiting factor. It is possible that a fold that can be built from a small amino acid alphabet is significantly smaller than or greater than the selected length.

Structure validation was successful for the LAGVSTDP composed of 8 amino acid types. We did not determine the full 3D structure of the protein using any structural biology method, but we are relatively confident with the predicted structure because it agrees with the ECD spectrum. This protein is a *β*-sheet-rich protein, which is generally recognized as challenging in protein design.

We successfully designed and experimentally verified the protein composed of 8 amino acid types. The predicted fibronectin type III domain architecture (supported by the shape of the ECD spectrum) is more complex than the helical bundle designed from 7 amino acid types (Schafmeister et al. (1997)). On the other hand, unlike many proteins designed by a reduction of amino acid alphabets of existing proteins (Riddle et al. (1997); Longo et al. (2013); Walter et al. (2005)) it was not designed to perform any enzymatic or binding function.

The designed LAGVSTDP protein belongs to a well-characterized fold. It was reported that `lm-design` sometimes designs 3D structures very similar to proteins with known 3D structures, and sometimes it hallucinates original structures (Verkuil et al. (2022)). In this case, the program reused the existing fold.

To our knowledge, the fibronectin type III domain is not recognized as an evolutionarily old fold. This indicates that also proteins that are not evolutionarily old can be redesigned in a reduced amino acid alphabet, or at least in the alphabet of size 8.

We were unable to produce the other two selected proteins from 6 and 4 amino acid types (LAGSID and LAID, respectively) in the amounts sufficient for experimental characterization. One possible explanation is that the designed proteins do not fold properly and were degraded or aggregated during the expression. This explanation is supported by low ProteinMPNN_sol_ confidence (0.16 and 0.14 for LAGSID and LAID, respectively) (Figure 5). The redesign of LAGSID and LAID resulted in the formation of long surface-exposed helices composed mainly of alanine. For this reason, we did not validate them experimentally.

Another explanation is that the expression failed for some biological reasons, for example that the cells run out of the corresponding aminoacyl-tRNAs, some secondary structures were formed on non-complex DNA or RNA, or others. Finally, it is possible that we produced the proteins but we were unable to purify them, as these proteins may exhibit unexpected behavior, such as poor staining on electrophoresis gels.

In the experimental tests of proteins designed by `lm-design`, the authors of this software reported a success rate of 67 % (Verkuil et al. (2022)). According to the binomial distribution, with the same success rate we can expect 22 % probability of successful production of one of the three designs. Therefore, our success rate does not contradict the result of the `lm-design` developers.

The fact that LAGVSTDP is poorly thermostable provides weak support for the model of its 3D structure. Inspired by the success of Giacobelli and co-workers (Giacobelli et al. (2025)) in using the ProteinMPNN family of programs, we redesigned LAGVSTDP by ProteinMPNN_sol_, keeping the same amino acid alphabet. The redesigned protein (LAGVSTDP2, with 69.7 % identity with LAGVSTDP) was stable up to 70 °C and, more importantly, its thermal denaturation was reversible. This significantly improves the confidence of the 3D model of LAGVSTDP and LAGVSTDP2.

In conclusion, we computationally demonstrated and experimentally verified that a stable fold can be formed of 8 amino acid types. An experimental validation is missing for smaller alphabets; nevertheless, we believe that we demonstrated that small alphabet proteins are interesting from an evolutionary as well as biotechnological point of view.

## 4 Methods

### 4.1 Computational methods

Analysis of reduced amino acid alphabets in modern proteins was performed using *ad hoc* Python scripts (see the Zenodo dataset). UniProt version 2025_3 was used (The UniProt Consortium (2024)).

ESMfold (ESM-2 Public Release v1.0.3, Lin et al. (2023)) and its `lm-design` modules (Verkuil et al. (2022)) were obtained from GitHub (https://github.com/facebookresearch/esm). The module was modified to enable work with reduced amino acid alphabets (by zeroing the probabilities of restricted amino acid types and renormalizing remaining ones, see the Zenodo dataset). For each amino acid alphabet, we performed a protein design run (170,000 Monte Carlo steps) starting from a random sequence. Simulated annealing was introduced as follows. The initial temperature was 8 and it was reduced to half every 10,000 steps. DSSP (version 4.4.7, Hekkelman et al. (2025)) was used to assign the secondary structure.

tSNE analysis (van der Maaten and Hinton (2008)) was performed using `scikit-learn`. The C-α atom coordinates were extracted from all the designed models and superimposed on the coordinates of the model designed for the alphabet of all early amino acids. The resulting coordinates (100 atoms, 300 coordinates *x, y*, and *z*) were used as input to tSNE with default settings.

The ProteinMPNN_sol_ model v48 (Goverde et al. (2024)) implemented using LigandMPNN software (Dauparas et al. (2025)) was used for the redesign of proteins. The sampling temperature was set to 0.1 (in *kT* units). We generated 50 sequences in each run and the one with the highest confidence was selected.

The values of pLDDT reported in this study refer to mean values for the C-α atoms of the models predicted by ESMfold (in %). `R, ggplot2` and the UCSF Chimera (Pettersen et al. (2004)) were used to make the figures. ANOVA was performed in the R software package. The identification of knots in proteins was performed using the topoly 1.1.0 library (Dabrowski-Tumanski et al. (2021)) using 2,000 trials of the Alexander polynomial routine.

Detailed datasets (modified `lm-design`, examples of raw design data, all resulting sequences and predicted 3D structure, data used to generate images, and detailed experimental procedures) are available via Zenodo (DOI: 10.5281/zenodo.18889431 and 10.5281/zenodo.22024387).

### 4.2 Experimental validation

The sequences of the designed proteins LAGVSTDP, LAGSID, and LAID were reverted to nucleotide sequences with codons chosen with probabilities based on the frequency distribution for expression in *E. coli* and synthesized as constructs in the pET15b expression vectors by GenScript Biotech, China. They were transformed into competent *E. coli* Lemo21(DE3) cells (New England Biolabs, USA) using a heat shock transformation. They were cultivated at 37 °C, 200 RPM in Luria-Bertani medium in the presence of ampicillin (100 mg/L). The expression of the protein was induced by the addition of isopropyl β-D-1-thiogalactopyranoside (0.3 mM) when the optical density reached OD_600nm_ = 0.5. The cells were further cultivated overnight at 22 °C. The cells were collected by centrifugation (8,000 g, 10 min, 4 °C), disrupted by One Shot Cell Disruptor (L&L Biosystems GmbH, Germany, pressure 2.7 kBar).

We used a similar protocol for the production of the LAGVSTDP2 with several differences. Namely, we used pMAL-c5X vector by GenScript Biotech and *E. coli* BL21(DE3) cells (New England Biolabs). After induction, the cells were further cultivated overnight at 16 °C.

To purify LAGVSTDP, LAGSID and LAID, the cell-free extract was loaded onto the NiNTA agarose column (Cube Biotech GmbH, Germany). The elution was monitored by SDS-PAGE (Laemmli (1970)). The elution was performed by an increasing concentration of imidazole (0-500 mM) in phosphate buffered saline (PBS, 10 mM Na_2_HPO_4_, 1.8 mM KH_2_PO_4_, 137 mM NaCl, 2.7 mM KCl, pH 7.4). Pooled fractions containing the expressed protein were desalted by gel filtration (PD-10 columns packed with Sephadex™ G-25 resin, Cytiva, USA, in PBS) and loaded onto a liquid chromatography system (NGC Discover 10 Chromatography System, BioRad, USA) equipped with the gel permeation chromatography column (HiPrep 16/60 Sephacryl™ S-100 HR, Cytiva). Due to the absence of aromatic amino acids in the designed proteins, the elution was monitored as absorbance at 210, 215, and 220 nm, in addition to the standard 280 nm, which was used to monitor possible impurities. PBS was used as a mobile phase. The elution was monitored by SDS-PAGE. The pooled fractions containing the desired protein were dialyzed against water (Spectra/Por 3, 3,500 Da cut-off, Repligen Corp., USA) and freeze-dried.

The cell-free extract containing LAGVSTDP2 was loaded onto MBPTrap™ HP column (Cytiva), washed with a PBS buffer (the same as above) and eluted with a PBS buffer containing 10 mM maltose. The pooled fractions containing the MBP-LAGVSTDP2 fusion protein (2 mL) were treated with factor Xa (New England Biolabs, diluted 1:25) overnight at room temperature. Next, the mixture was loaded onto a gel permeation chromatography column as described above. The elution was monitored as absorbance at 205 nm, in addition to the standard 280 nm. The pooled fractions containing the desired protein were dialyzed against water (Spectra/Por 3, 3,500 Da cut-off, Repligen Corp.) and freeze-dried.

The freeze-dried samples were reconstituted in 25 mM phosphate buffer (K_2_HPO_4_/KH_2_PO_4_, pH 7.5) to reach the concentration of 0.20 mg/mL (measured by the Bicinchoninic acid method, Pierce™ BCA Protein Assay Kit, Thermo Fisher Scientific Inc., USA). ECD and UV absorption spectra were collected using the J-815 spectrometer (Jasco International Co., Ltd., Japan) in a 1 mm quartz cuvette (Hellma GmbH & Co. KG, Germany) at a 50 nm/min scanning speed, 2 s response time, and a bandwidth of 2 nm in the 180-260 nm region. The spectrometer was purged during measurements by gaseous nitrogen (purity > 99.99 %, SIAD Czech spol. s r.o., Czech Republic). See the Zenodo dataset for the reference UV absorption spectrum. The final spectra were presented as an average of six spectral accumulations. The baseline correction was performed by subtracting the spectrum of the buffer solution measured under identical experimental conditions. The temperature was controlled with a Jasco Peltier CDF-426S/16 temperature control system (Jasco International Co., Ltd.). The corresponding absorbance values were calculated from the photomultiplier HT voltage values using the Spectra analysis module of Spectra Manager (Jasco International Co., Ltd., see the Zenodo dataset).

## 5 Acknowledgment

We thank Tereza Neuwirthová, Filip Buchel, Valerio Guido Giacobelli, and Klára Hlouchová for a fruitful discussion. The project was supported by COST (project ML4NGP, CA21160). Participation in the COST project was supported by the Ministry of Education, Youth, and Sports of the Czech Republic (LUC 24136). Computation was supported by ELIXIR CZ and “e-Infrastruktura CZ” (LM 2023055 and e-INFRA CZ ID: 90254).

